# Sex-specific Effects of the Endocannabinoid Agonist 2-Arachidonoylglycerol on Sleep and Circadian Disruptions during Fentanyl Withdrawal

**DOI:** 10.1101/2023.12.19.572466

**Authors:** Mackenzie C. Gamble, Sophia Miracle, Benjamin R. Williams, Ryan W. Logan

**Author notes:** **Corresponding Author:** Ryan W. Logan, PhD Professor Department of Psychiatry Department of Neurobiology University of Massachusetts Chan Medical School LRB, Office 611 364 Plantation Street, Worcester, MA 01605.

## Abstract

Fentanyl has become the leading driver of opioid overdoses. Cessation of opioid use represents a challenge as the experience of withdrawal drives subsequent relapse. One of the most prominent withdrawal symptoms that can contribute to opioid craving and vulnerability to relapse is sleep disruption. The endocannabinoid agonist, 2-Arachidonoylglycerol (2-AG), may promote sleep and reduce withdrawal severity; however, the effects of 2-AG on sleep disruption during opioid withdrawal have yet to be assessed. Here, we investigate the effects of 2-AG administration on sleep-wake behavior and diurnal activity in mice during withdrawal from fentanyl. Sleep-wake activity was continuously recorded before and after chronic fentanyl administration in both male and female C57BL/6J mice. Immediately following cessation of fentanyl administration, 2-AG was administered intraperitoneally to investigate the impact of endocannabinoid agonism on opioid-induced sleep disruption. Female mice maintained higher activity levels in response to chronic fentanyl than male mice. Furthermore, fentanyl increased wake and decreased sleep during the light period and inversely increased sleep and decreased wake in the dark period in both sexes. 2-AG treatment increased arousal and decreased sleep in both sexes during first 24 hrs of withdrawal. On withdrawal day 2, only female showed increased wakefulness with no changes in males, but by withdrawal day 3 male mice displayed decreased rapid-eye movement sleep during the dark period with no changes in female mice. Overall, repeated administration of fentanyl altered sleep and diurnal activity and administration of the endocannabinoid agonist, 2-AG, had sex-specific effects on fentanyl-induced sleep and diurnal changes.

## 1. Introduction

Opioid use disorder (OUD) remains a major public health concern. Currently, synthetic opioids are responsible for the majority of opioid overdose deaths, with the proliferation of fentanyl driving the opioid epidemic (Ahmad FB et al., 2023). Fentanyl is the most commonly misused synthetic opioid, with higher efficacy, quicker onset, and greater potency than most other opioids (Bird et al., 2023). Opioid use can lead to dependence and subsequent withdrawal which is associated with a number of negative symptoms (Shah and Huecker, 2023). This includes sleep disturbances which occur during withdrawal and into protracted abstinence (Kay, 1975; Lewis et al., 1970; Oswald, 1969). This lack of sleep, in turn, can drive relapse. Few studies have interrogated the impact of opioid withdrawal on sleep and discovering new treatments that mitigate sleep disruptions during withdrawal is a growing area of interest. Notably, targets within the endocannabinoid (eCB) system pose potential in attenuating opioid withdrawal symptoms (Galaj and Xi, 2019).

Cannabinoid receptor 1 and 2 (CB1/2R) agonists, such as dronabinol, may reduce overall opioid withdrawal severity with evidence in both rats and humans (Sloan et al., 2017). Indeed, the eCB system has been implicated in reward and reinforcement of opioids (for review see Wenzel and Cheer, 2018). For example, opioid and eCBs synergistically modulate each other. Δ^9^-tetrahydrocannabinol administration increases endogenous opioid levels in the nucleus accumbens (NAc) while opioid administration in turn increases eCB levels in the NAc (Caillé et al., 2007; Valverde et al., 2001). Moreover, CB1 knockout blocks development of conditioned place preference and self-administration in rodents (Martin et al., 2000; Navarro et al., 2004). CB1R and mu-opioid receptor co-localize in the NAc core and shell, particularly GABA neurons, (Pickel et al., 2004) which are central to opioid reinforcement and withdrawal (Chieng and Williams, 1998; Saigusa et al., 2021; Xi and Stein, 2002). 2-Arachidonoylglycerol (2-AG), is a full agonist of both the CB1R and CB2R (with greater affinity for the CB1R) (Kendall and Yudowski, 2017) abundant in the central nervous system including reward circuitry (Sugiura et al., 2006). Acute fentanyl administration, chronic subcutaneous morphine injections, and heroin self-administration have all shown to decrease 2-AG levels in the NAc of rats (Caillé et al., 2007; Sustkova-Fiserova et al., 2017; Viganò et al., 2003). Additionally, 2-AG administered intracerebrally or increased by global inhibition of its primary degradation enzyme, monoacylglycerol lipase (MAGL), reduced behavioral withdrawal symptoms in morphine dependent mice both following naloxone-precipitated (Ramesh et al., 2011; Yamaguchi et al., 2001) as well as spontaneous withdrawal (Ramesh et al., 2013).

Beyond addiction, the eCB system including 2-AG has been associated with sleep-wake regulation (for review see Kesner and Lovinger, 2020). Bilateral intracerebral administration of 2-AG was shown to increase REMS in rats (Pérez-Morales et al., 2013) while increasing 2-AG tone via inhibition of MAGL mostly increased NREMS when administered before the dark period and mainly decreased REMS when administered before the light period in mice (Pava et al., 2016). Evidence from blood serum taken from humans indicates that 2-AG has a diurnal rhythm with a nadir in the middle of the sleep period which is enhanced under sleep restriction (Hanlon et al., 2016). Furthermore, CB1 inverse agonist, rimonabant, has been noted to induce insomnia in some individuals (Nathan et al., 2011) and CB1 knockout mice have increased wake and decreased NREMS (Pava et al., 2014). While evidence suggests that the endocannabinoid system including 2-AG plays a role in addiction and sleep, no studies to date have looked at how it may affect withdrawal-induced sleep disruption. Thus, we investigated the impact of 2-AG administration at the onset of withdrawal following chronic fentanyl administration on sleep-wake behavior.

## 2. Methods

### 2.1. Animals and Housing

Adult (10-15 weeks old) male and female C57BL/6J mice (24 total, balanced by sex) from Jackson Laboratory (Strain #000664) were used. Upon arrival, mice were grouped housed in a standard 12:12 light dark cycle (lights on at 0700h and off at 1900h) with rodent chow and water provided *ad libitum.* For sleep-wake actigraphy, mice were single housed in sleep boxes within a sound attenuating light controlled ventilated cabinet. Animals were habituated for at least three days before any recording. Cage floors were cleaned every four days to limit disruptions to sleep and activity recordings. All experimental procedures were approved by the Institutional Animal Care and Use Committee at Boston University School of Medicine.

### 2.2. Drugs

Fentanyl citrate (50 mg/mL) was dissolved in 0.9% sterile filtered saline to reach a final concentration of 32 μg/mL. 2-Arachidonoylglycerol (2-AG) was obtained from Tocris Biosciences (Cat. number: 1298/10) and stored at -80C. The day before administration 2-AG was dissolved in 10% dimethyl sulfide (DMSO) diluted with sterile saline at a final concentration of 1 mg/mL and dosed at 5 mg/kg.

### 2.3. Chronic Fentanyl Exposure and 2-Arachidonoylglycerol Administration

Mice were habituated to the sleep boxes for three days, after which baseline wake and sleep activity was recorded (day 1). Mice underwent chronic fentanyl administration to induce physical dependence (Neumueller et al., 2023; Puig et al., 2022). Mice received twice daily intraperitoneal (i.p.) injections of fentanyl at 320 μg/kg for 7 days (day 2-8) (Gamble et al., 2022; Puig et al., 2022). Injections were approximately 8 hours apart each day (0730h then 1530h). From withdrawal day 1 (WD1) to WD3, mice received once daily i.p. injection of either 2-AG or vehicle (10% DMSO) at 5 mg/kg (0730h).

### 2.4. Actigraphy Recording

Throughout the experiment, mice were housed in individual 6-inch x 6-inch polycarbonate cages (Signal Solutions: Sleep Monitoring System for Mice) set with floor sensor pads that were attached to a recording laptop. Activity signals were recorded, identified, and categorized as Wake, NREMS, or REMS using SleepStats software version 4 (Signal Solutions), binned in 30-minute intervals. All wake, NREMS, and REMS data are presented normalized to each mouse’s own post-habituation baseline recording. Thus, all sleep measures represent a change from baseline. Activity data was further exported as 1 min bins into ClockLabs Analysis 6 software (Actimetrics) to generate the diurnal activity plots.

### 2.5. Statistics

All data was analyzed as two-way repeated measures ANOVA using GraphPad Prism (version 9.4.1). Sidak post-hoc corrections were used for multiple comparisons when interaction effects were significant. Results are presented as mean ± standard error of the mean (SEM) for data plotted by Zeitgeber Time (ZT) or light-dark period, and 95% confidence intervals when collapsed by day with significance set at α□=□0.05.

## 3. Results

### 3.1. Female mice displayed greater activity in the dark portion of the light-dark cycle in response to chronic fentanyl relative to males

______Mice were administered fentanyl (320 μg/kg) twice daily eight hours apart for seven days and activity averaged across all seven days of administration (Fig. 1). A significant interaction effect between sex and time was found (F_(23, 506)_ = 5.821, P<0.0001) in mean activity duration. A main effect was observed for time (F_(7.609, 167.4)_ = 101.0, P<0.0001), but not sex (F_(1, 22)_ = 3.882, P=0.0615). In both male and female mice, activity increased in response to to each fentanyl injection, likely concordant with handling and opioid-induced locomotor sensitization (Fig. 2). During this period, post-hoc analysis revealed that in males the second daily fentanyl injection induced greater activity at ZT10 than females (P=0.0024). Another peak of activity was seen at the onset of dark phase (active period) in both sexes (Fig. 2). Following this peak, at ZT16, 17, and 18, female mice had increased activity compared to males (P = 0.0073, P = 0.0238, P = 0.0041).

**Fig. 1:**
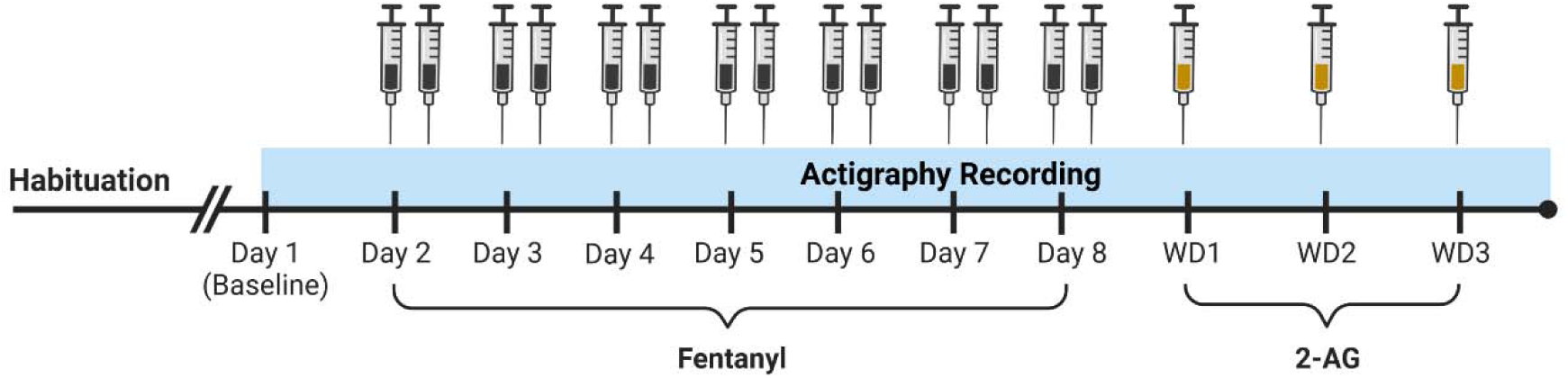
Schematic of experimental design. Animals were habituated to the recording chamber before a pre-drug baseline was established (at least 3 days). All animals received fentanyl twice daily for seven days. At the onset of withdrawal half of the mice received 2-AG treatment or vehicle (10% DMSO) once daily for three days.

**Fig. 2:**
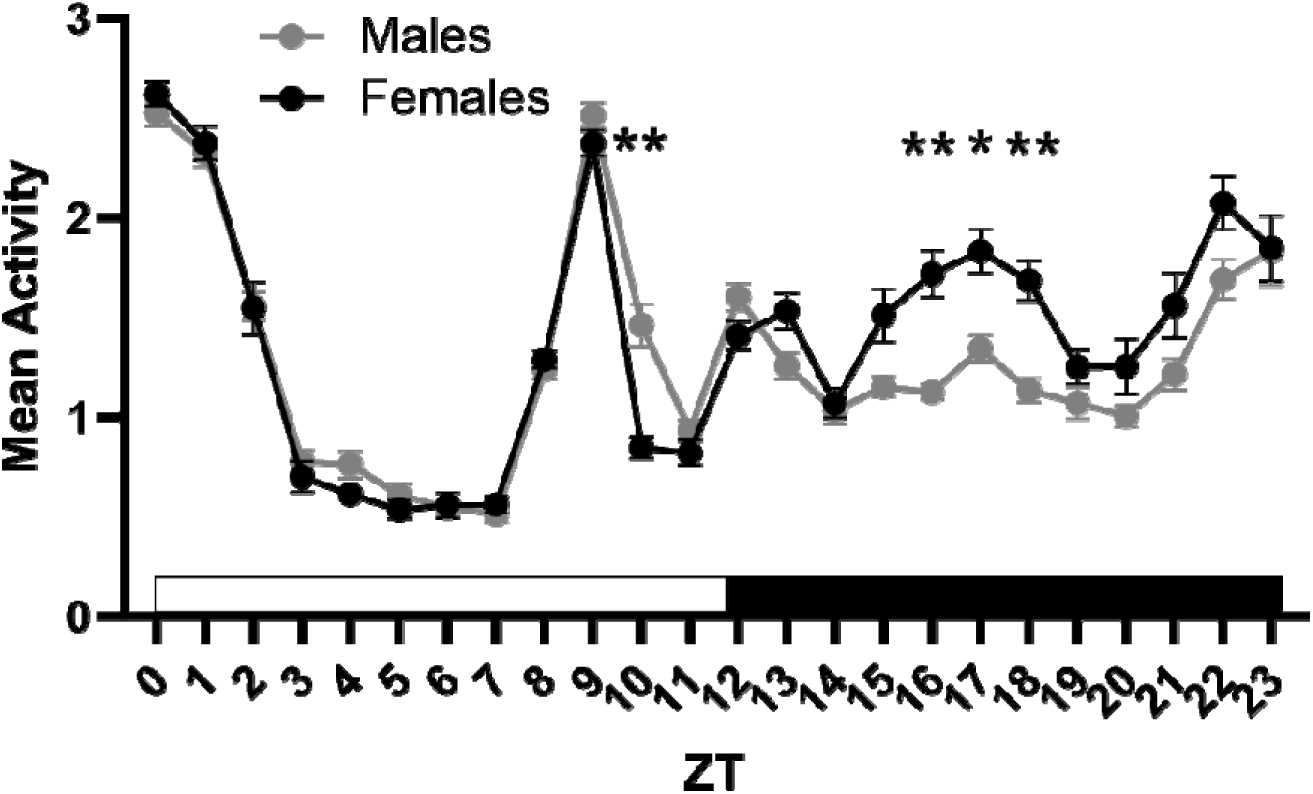
Diurnal activity between males and females averaged in 6-minute bins per ZT across all 7 days of fentanyl administration. Data presented as mean ± SEM. * = P ≤ 0.05, ** = P ≤ 0.01

### 3.2. Chronic fentanyl administration increased NREMS and REMS in males and females

Sleep-wake behavior was recorded using actigraphy, normalized to an undisturbed baseline recording preceding fentanyl. Each fentanyl injection immediately increased wake above baseline in both sexes, lasting ∼2-4 hours (Fig. 3a). Following this 2–4-hour period of increased wake after the first injection there was minimal change in sleep-wake from baseline, except on day 1 and to a lesser extent day 2 where REMS is perturbed in opposing directions with females showing an increase and males a decrease. Following the 2-4 hours from the second injection which coincides with the onset of the dark period (active period) there was a large depression of wake and an increase in both NREMS and REMS for the rest of the period in both sexes (Fig. 3b and c).

**Fig. 3:**
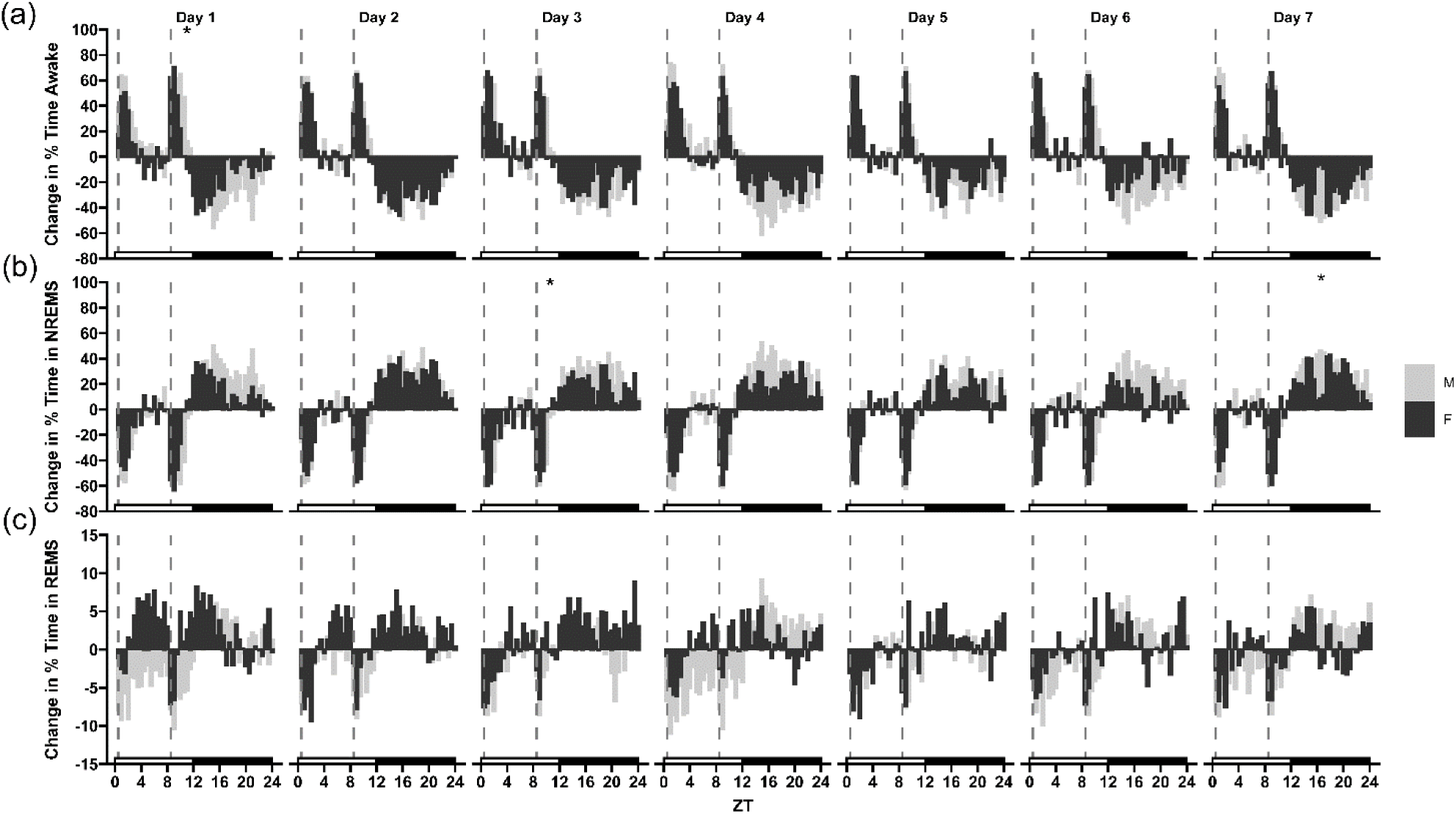
Change in the percent time in wake, NREMS, and REMS from baseline during chronic fentanyl administration for male and female mice. The data are presented as means per half hour (SEM omitted for clarity) with vertical dashed lines indicating daily fentanyl injections. * = P ≤ 0.05

A significant interaction effect between sex and time on the change in percent time awake was found on days 1 (F_(47, 1034)_ = 2.308, P<0.0001), 4 (F_(47, 1034)_ = 1.844, P=0.0006), 6 (F_(47, 1034)_ = 1.867, P=0.0004), and 7 (F_(47, 1034)_ = 2.161, P<0.0001). A main effect of sex was seen only on day 6 (F_(1, 22)_ = 8.173, P=0.0091) and time on all corresponding interaction effect days (Day 1: F_(11.90, 261.8)_ = 19.16, P<0.0001; Day 4: F_(13.20, 290.4)_ = 22.11, P<0.0001; Day 6: F_(11.18, 245.9)_ = 14.08, P<0.0001; Day 7: F_(14.29, 314.4)_ = 20.70, P<0.0001). Similarly, there was a significant interaction effect (sex x time) on the change in percent time in NREMS on days 1 (F_(47, 1034)_ = 1.870, P=0.0004), 3 (F_(47, 1034)_ = 1.413, P=0.0366), 4 (F_(47, 1034)_ = 1.447, P=0.0277), 6 (F_(47, 1034)_ = 1.737, P=0.0018), and 7 (F_(47, 1034)_ = 2.379, P<0.0001) with a significant main effect of sex only on days 4 (F_(1, 22)_ = 6.665, P=0.0170) and 6 (F_(1, 22)_ = 12.20, P=0.0021). A significant main effect of time was found for all corresponding interaction effect days (Day 1: F_(12.33, 271.3)_ = 19.46, P<0.0001; Day 3: F_(13.34, 293.6)_ = 18.86, P<0.0001; Day 4: F_(13.29, 292.3)_ = 21.64, P<0.0001; Day 6: F_(12.22, 268.8)_ = 14.37, P<0.0001; Day 7: F_(14.49, 318.8)_ = 20.71, P<0.0001). Lastly, there was a significant interaction effect on the change in percent time in REMS on days 1 (F_(47, 1034)_ = 2.619, P<0.0001), 4 (F_(47, 1034)_ = 3.095, P<0.0001), 6 (F_(47, 1034)_ = 1.495, P=0.0182), and 7 (F_(47, 1034)_ = 1.685, P=0.0030) as well as a main effect of sex on day 3 (F_(1, 22)_ = 4.441, P=0.0467) only. A significant main effect of time was present for all corresponding interaction effect days (Day 1: F_(10.34, 227.4)_ = 3.996, P<0.0001; Day 3: F_(11.94, 262.6)_ = 4.118, P<0.0001; Day 4: F_(11.94,262.8)_ = 6.110, P<0.0001; Day 6: F_(10.88, 239.3)_ = 3.874, P<0.0001; Day 7: F_(11.33, 249.2_) = 5.454; P<0.0001). Post-hoc analysis (Sidak corrected) revealed that ZT 10.5 (P= 0.0423) of day 1 for wake was significantly higher in males compared to females. For NREMS, ZT 10 (P=0.0399) of day 3 was significantly lower in males compared to females and ZT 16 (P=0.0449) of day 7 was significantly higher in males compared to females (Fig. 3a and b). Ultimately, wakefulness increased and sleep decreased during the light period while the inverse occurred in the dark period in both sexes compared to individual baselines across all days of fentanyl administration.

Both males and females had a significant main effect of time (Female: F_(11.98, 263.5)_ = 15.38, P<0.0001; Male: F_(12.28, 270.2)_ = 26.04, P<0.0001) and a significant interaction effect between time and change in percent time awake from baseline on day 1 compared to day 7 (Female: F_(47, 1034)_ = 1.472, P=0.0223; Male: F_(47, 1034)_ = 1.605, P=0.0067). Post-hoc analysis indicated no ZTs were significantly changed in females between day 1 and day 7 of fentanyl administration, but in males ZT 10 was significantly elevated (P=0.0265). Similarly, there was a significant main effect of time for females and males (Female: F_(11.87, 261.2)_ = 16.69, P<0.0001; Male: F_(12.50, 274.9)_ = 24.23, P<0.0001) and day for females (F_(1, 22)_ = 4.318, P=0.0496) for NREMS. Both sexes had a significant interaction effect (Female: F_(47, 1034)_ = 1.500,P=0.0175; Male: F_(47, 1034)_ = 1.782, P=0.0011), but only males had a significant change from day 1 to day 7 following post-hoc analysis with a decrease in NREMS at ZT 10 (P=0.0187). No change was found for REMS between day 1 and day 7. In summary, ZT 10 males displayed about 50% less wake (and 45% more NREMS) by day 7 of fentanyl administration. Otherwise changes in sleep-wake state remained largely the same between first and last fentanyl injection.

### 3.3. 2-AG did not alter diurnal activity in male or female mice during withdrawal

Following chronic fentanyl administration, mice underwent spontaneous withdrawal and received once daily injections of vehicle (DMSO) or 2-AG at the onset of the light period each day for 3 days of withdrawal (Fig. 4). Neither DMSO nor 2-AG had any significant effect on activity across the light or dark phase in both males (Treatment: F_(1, 10)_ = 2.299, P=0.1604, Time: F_(4.108, 41.08)_ = 35.53, P<0.0001, Treatment x Time: F_(23, 230)_ = 0.8047, P=0.7241) and females (Treatment: F_(1, 10)_ = 4.411, P=0.0620, Time: F_(5.567, 55.67)_ = 37.17, P<0.0001, Treatment x Time: F_(23, 230)_ = 1.329, P=0.1499) during withdrawal (Fig. 4a and b).

**Fig 4:**
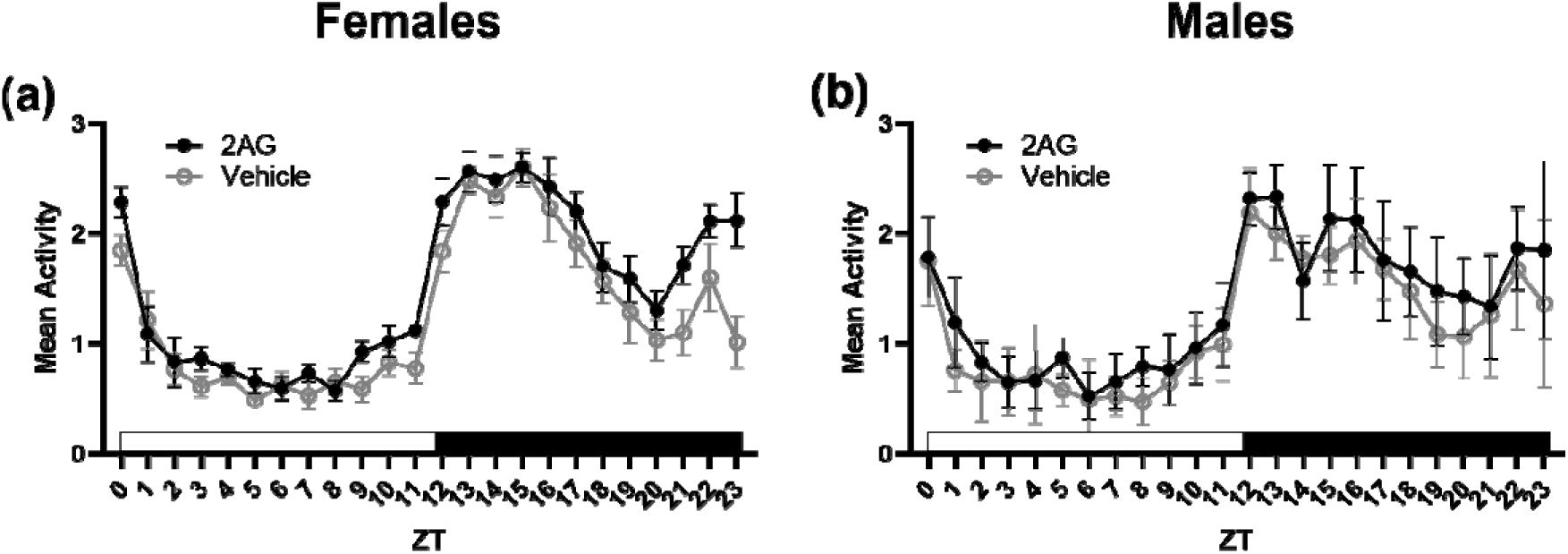
Diurnal activity between male and female mice averaged in 6 min bins per ZT across all 3 days of withdrawal. Data presented as mean ± SEM.

### 3.4. 2-AG increased wakefulness in both sexes and decreased REMS in female mice during withdrawal day 1

A significant main effect of treatment and time on the change in percent time awake was found for withdrawal day 1 in females (F_(1, 10)_ = 11.41, P < 0.0013; F_(5.578, 55.78)_ = 4.405, P < 0.0070). 2-AG treated female mice showed increased in wake compared to vehicle (DMSO) controls across the 24-hour withdrawal period (8.76[2.00,15.52] vs -7.37[- 13.80,0.923]). Change in percent time awake was above baseline levels in 2-AG treated mice and below baseline levels in vehicle controls (Fig. 5a). Furthermore, there was a significant interaction effect of treatment and time (F_(47, 470)_ = 2.277, P <0.0001) and multiple comparisons analysis indicated 2-AG treatment significantly increased the change in percent time awake relative to vehicle controls at ZTs 3.5 (14.11 ± 3.40 vs -42.43 ± 9.16, P = 0.0448), 12 (51.96 ± 8.17 vs -15.58 ± 3.142, P = 0.0084), 12.5 (37.46 ± 5.49 vs -14.41 ± 7.66, P = 0.0176), and 24 (-3.07 ± 3.18 vs -24.3 ± 1.97, P = 0.0192) in females (Fig. 5a). 2-AG also significantly increased the change in percent time wake compared to vehicle controls (-0.8647[-5.648,3.919] vs -10.67[-16.20,5.134]) across the 24-hour withdrawal period in males (Treatment: F_(1, 10)_ = 14.42, P=0.0035, Time: F_(4.713, 47.13)_ = 2.682, P=0.0352). As opposed to females, levels of wake in males were on par with baseline levels in 2-AG treated mice. Vehicle controls as was observed in females were below baseline levels (Fig. 5b). Post-hoc analysis following a significant interaction effect (F_(47, 470)_ = 2.366, P<0.0001) indicated that 2-AG treatment significantly increased the change in percent time awake relative to vehicle controls at ZTs 2.5 (27.85±8.37 vs - 26.2±4.33, P = 0.0263) and 20 (20.61±9.48 vs -34.76± 5.36, P = 0.0462) in males (Fig. 5b). Change in average wake bout duration had a significant main effect of treatment (Females: F_(1, 10)_ = 7.312, P=0.0222; Males: F_(1, 10)_ = 11.80, P=0.0064) as well as a significant interaction effect in both sexes (Females: F_(1, 10)_ = 5.320, P=0.0438; Males: F_(1, 10)_ = 18.40, P=0.0016). Multiple comparisons revealed that wake bouts were longer in 2-AG treated than vehicle controls, but only in the dark period (Females: 133.82 ± 21.26 vs - 200.33 ± 131.57, P = 0.004, Males: 40.86±36.812 vs -141.72±19.00, P < 0.0001) (Fig. 5c and d). 2-AG increased the change in average wake duration above baseline while vehicle decreased the change in average wake duration below baseline in the dark period.

**Fig. 5:**
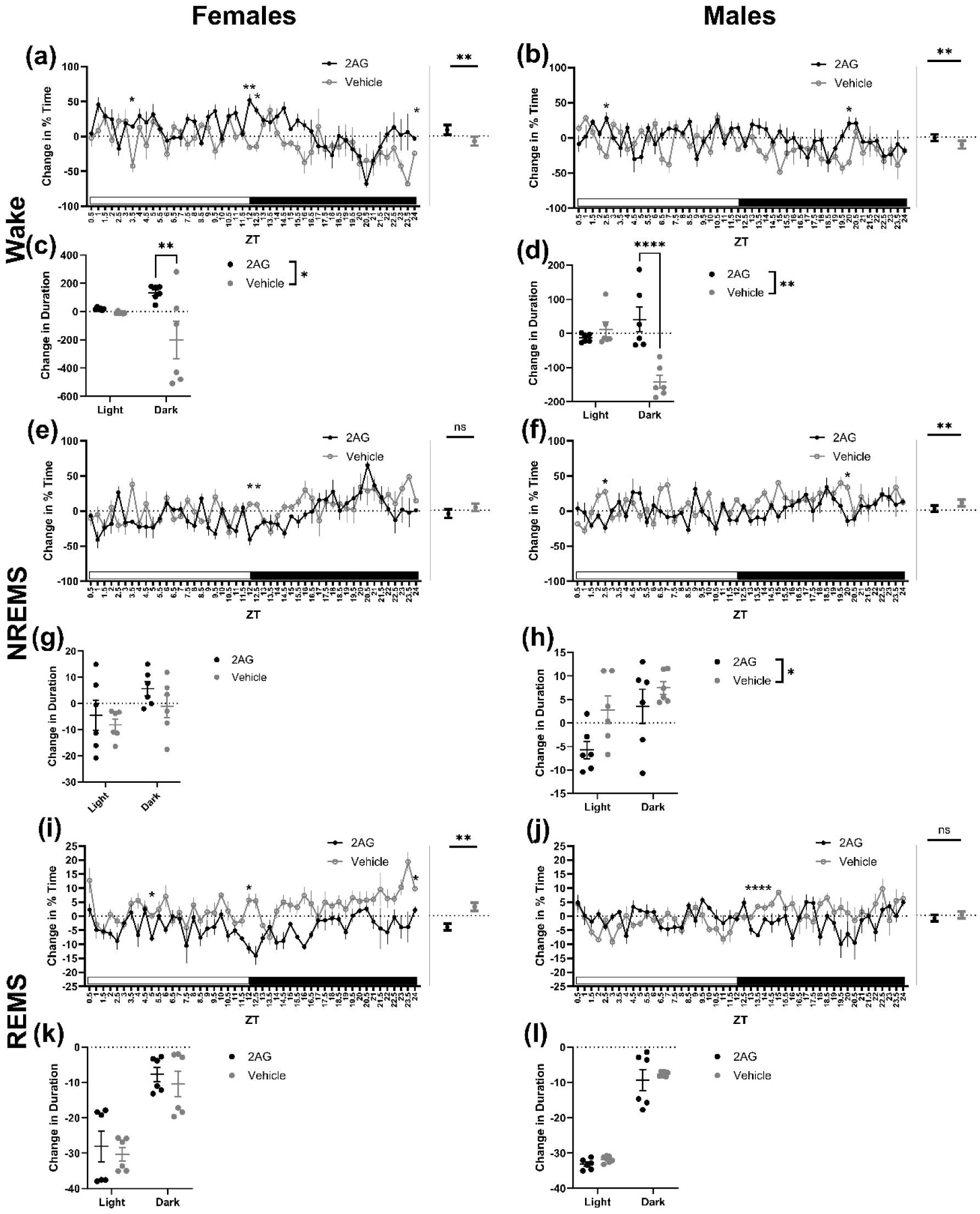
Sleep-wake changes in male and female mice during withdrawal day 1. (a) Half hourly (left) and total (right) change in percent time spent awake from baseline in females. (b) Half hourly (left) and total (right) change in percent time spent awake from baseline in males. (c) Change in wake bout duration from baseline in females during the light and dark period. (d) Change in wake bout duration from baseline in males during the light and dark period. (e) Half hourly (left) and total (right) change in percent time in NREMS from baseline in females. (f) Half hourly (left) and total (right) change in percent time in NREMS from baseline in males. (g) Change in NREMS bout duration from baseline in females during the light and dark period. (h) Change in NREMS bout duration from baseline in males during the light and dark period. (i) Half hourly (left) and total (right) change in percent time in NREMS from baseline in females. (j) Half hourly (left) and total (right) change in percent time in NREMS from baseline in males. (k) Change in REMS bout duration from baseline in females during the light and dark period. (l) Change in REMS bout duration from baseline in males during the light and dark period. Data presented as mean ± SEM for half hourly data and mean ± 95% CI for total data. ns = P > 0.05, * = P ≤ 0.05, ** = P ≤ 0.01, *** = P ≤ 0.001

Female mice did not have a significant main effect of treatment (F_(1, 10)_ = 4.623, P = 0.0571). only time (F_(5.705, 57.05)_ = 4.234, P=0.0016) on the change in percent time in NREMS. Across 24 hours, 2-AG treated female mice were slightly below baseline levels while vehicle controls were slightly above baseline levels. A significant interaction effect (F_(47, 470)_ = 2.282, P<0.0001) followed by post-hoc analysis revealed that 2-AG treated female mice had significantly less NREMS at the transition from light to dark including ZT 12 (-40.555±8.26 vs 9.89±3.36, P = 0.0440) and ZT 12.5 (P = 0.0287) (Fig. 5e). Male mice, however, had a significant main effect of treatment and time (F_(1, 10)_ = 17.42, P=0.0019; F_(4.943, 49.43)_ = 2.632, P=0.0351). Change in percent time in NREMS was significantly decreased in 2-AG treated male mice compared to vehicle controls across 24 hours (1.878 [-2.488, 6.244] vs 10.48 [5.543, 15.42]). Vehicle controls had elevated NREMS compared to baseline while 2-AG treated mice had NREMS levels at baseline. A significant interaction effect (F_(47, 470)_ = 2.269, P<0.0001) followed by post-hoc analysis indicated that 2-AG treated male mice had decreased change in percent time in NREMS at ZTs 2.5 (-24.08 ± 7.81 vs 27.56 ± 4.52, P = 0.0209) and 20 (-14.26 ± 8.03 vs 33.61 ± 5.18, P = 0.0400) (Fig. 5f). NREMS bout durations were significantly lower in 2-AG treated male mice compared to vehicle controls (-1.145 ± 4.624 vs 5.139 ± 2.350; F_(1, 10)_ = 5.033, P=0.0487) with no significant interaction effect (F_(1, 10)_ = 0.8743, P=0.3718) (Fig. 5h). No differences in NREMS bout durations were found in females (Fig. 5g).

In female mice, there were significant main effects of treatment (F_(1, 10)_ = 17.32, P=0.0019) and time (F_(4.353, 43.53)_ = 3.903, P=0.0071) on the change in the percent time in REMS. 2-AG treatment significantly decreased REMS compared to vehicle controls across 24 hours (-4.058 [-5.258, -2.857] vs 3.235 [1.796, 4.675]) pushing levels below baseline while vehicle was notably above baseline. Moreover, post-hoc analysis following a significant interaction effect (F_(47, 470)_ = 2.348, P<0.0001) revealed change in percent time in REMS was significantly lower in 2-AG treated female mice at ZTs 5 (-7.97 ± 1.027 vs 0.11 ± 1.33, P = 0.0395), 12 (-11.41 ± 1.95 vs 5.69 ± 2.29, P = 0.0107), and 24 (2.20 ± 1.20 vs 9.80 ± 0.99, P = 0.0331) compared to vehicle controls (Fig. 5i). In males, the main effect of treatment was not significant (F_(1, 10)_ = 0.7257, P=0.4142) only time (F_(4.472, 44.72)_ = 2.538, P=0.0472). An interaction effect of treatment and time was found (F_(47, 470)_ = 3.053, P<0.0001) and post-hoc analysis revealed that ZT 13.5 (-6.775 ± 0.40 vs 3.48 ± 0.47, P<0.0001) had significantly decreased NREMS in 2-AG treated male mice compared to vehicle controls (Fig. 5j). No differences in REMS bout durations were found for either sex (Fig. 5k and l).

### 3.5. 2-AG increased wakefulness and decreased NREMS in females during withdrawal day 2

2-AG treatment in females had a significant main effect of treatment (F_(1, 10)_ = 7.574, P=0.0204) on the change in percent time awake while males did not (F_(1, 10)_ = 1.544, P=0.2424). Neither had a significant main effect of time. In females, wake was significantly elevated compared to vehicle controls with levels slightly above baseline while vehicle controls were slightly below (5.396 [0.8308, 9.960] vs -6.106 [-12.00, -0.2122]). Both sexes had a significant interaction effect (Females: F_(47, 470)_ = 2.365, P<0.0001; Males: F_(47, 470)_ = 2.639, P<0.0001). Wake was depressed at ZT 6.5 (-25.09 ± 2.60 vs 8.61 ± 4.87, P = 0.0165) in 2-AG treated females despite overall levels significantly elevated across 24 hours following post-hoc analysis (Fig. 6a). While in 2-AG treated males post-hoc analysis showed that wake was significantly elevated at ZT 5 (36.04 ± 10.64 vs -36.31 ± 9.73, P=0.0254) and then depressed at ZT 6 (-52.48 ± 5.43 vs 12.74 ± 3.95, P = 0.0002) (Fig 6b). Moreover, change in wake bout duration was significantly increased in 2-AG treated females across the entire second withdrawal day (43.58 ± 50.50 vs -30.60 ± 2.57, F_(1, 10)_ = 6.233, P=0.0316), but no effect was seen in males (Fig. 6c and d).

**Fig. 6:**
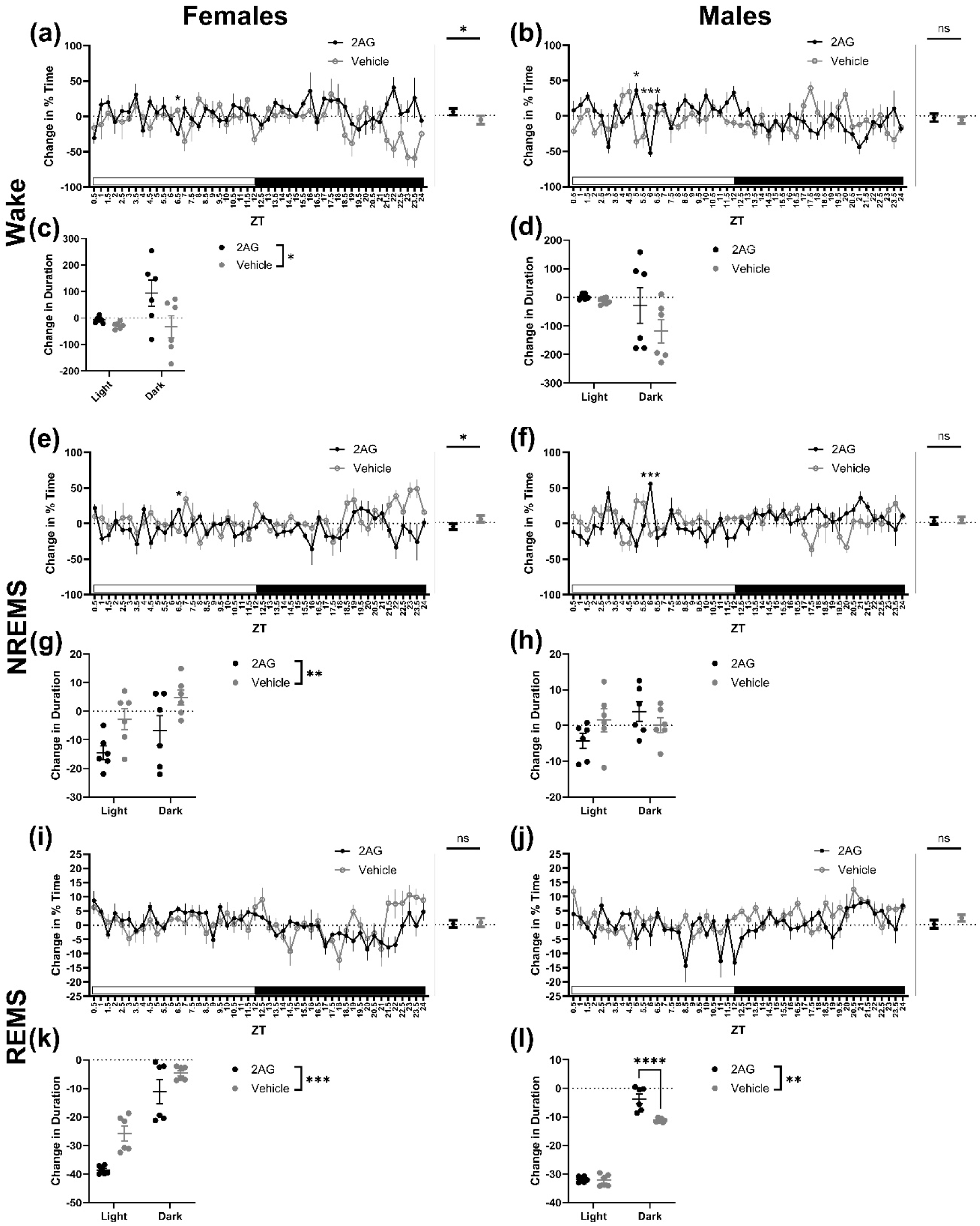
Sleep-wake changes in male and female mice during withdrawal day 2. (a) Half hourly (left) and total (right) change in percent time spent awake from baseline in females. (b) Half hourly (left) and total (right) change in percent time spent awake from baseline in males. (c) Change in wake bout duration from baseline in females during the light and dark period. (d) Change in wake bout duration from baseline in males during the light and dark period. (e) Half hourly (left) and total (right) change in percent time in NREMS from baseline in females. (f) Half hourly (left) and total (right) change in percent time in NREMS from baseline in males. (g) Change in NREMS bout duration from baseline in females during the light and dark period. (h) Change in NREMS bout duration from baseline in males during the light and dark period. (i) Half hourly (left) and total (right) change in percent time in NREMS from baseline in females. (j) Half hourly (left) and total (right) change in percent time in NREMS from baseline in males. (k) Change in REMS bout duration from baseline in females during the light and dark period. (l) Change in REMS bout duration from baseline in males during the light and dark period. Data presented as mean ± SEM for half hourly data and mean ± 95% CI for total data. ns = P > 0.05, * = P ≤ 0.05, ** = P ≤ 0.01, *** = P ≤ 0.001.

A significant main effect of treatment on the change in percent time in NREMS was found in females (F_(1, 10)_ = 9.734, P=0.0109), but not males (F_(1, 10)_ = 0.2312, P=0.6410). Neither had a significant main effect of time. As opposed to wake, NREMS in 2-AG treated female mice was significantly decreased compared to vehicle controls (-5.688 [-9.992, - 1.385] vs 5.315 [0.3236,10.31]). 2-AG treatment was slightly below baseline and vehicle controls slightly above baseline. Both sexes had a significant interaction effect (Females: F(_47, 470)_ = 2.330, P<0.0001; Males: F_(1, 10)_ = 0.2312, P=0.6410) with NREMS elevated at ZT 6.5 (19.46 ± 2.86 vs -10.98 ± 4.18, P = 0.0102) in females and ZT 6 (55.98 ± 3.01 vs -15.03 ± 5.56, P = 0.0102) in males following post-hoc analysis (Fig 6e and f). Again, as opposed to wake, change in NREMS bout duration significantly decreased across the entire second withdrawal day in females only (-10.64 ± 3.844 vs 1.043 ± 3.803, F_(1, 10)_ = 10.45, P=0.0090) (Fig. 6g).

No significant differences in the change in the percent time of REMS were found between 2-AG treated and vehicle control mice for either sex (Fig. 6i and j). Differences in the change in REMS duration, however, were seen in both sexes with a main effect of treatment (Females: F_(1, 10)_ = 23.33, P=0.0007; Males: F_(1, 10)_ = 14.3, P=0.0036) and time (Females: F_(1, 10)_ = 68.95, P<0.0001; Males: F_(1, 10)_ = 769.8, P = P<0.0001). In both sexes, 2-AG treated mice showed a significant reduction in the change in REMS bout duration compared to vehicle controls (Females: -24.83 ± 13.75 vs -15.23 ± 10.57; Males: -17.79 ± 14.10 vs -21.71 ± 10.53). Post-hoc analysis following a significant interaction effect in males (F_(1, 10)_ = 16.20, P=0.0024) indicated that this effect was restricted to the dark period (-3.68 ± 1.66 vs -11.18 ± 0.29, P<0.0001) (Fig. 6k). No interaction effect was found in females (Fig. 6l)

### 3.6. 2-AG disrupted REMS in only male mice during withdrawal day 3

By withdrawal day 3, no differences between 2-AG treated and vehicle control (main effect of treatment) female mice were evident in percent time in wake or sleep with most levels comparable to baseline across 24 hours. In male mice, there was a significant interaction effect on both the change in percent time awake (F_(47, 470)_ = 2.287, P<0.0001) and in NREMS (F_(47, 470)_ = 2.240, P<0.0001), but not treatment. Post-hoc analysis indicated 2-AG treatment simultaneously increased wake (18.53 ± 7.27 vs -24.72 ± 4.91, P=0.0411) and decreased NREMS (-17.665 ± 6.77 vs 22.42 ± 5.08, P=0.0460) at ZT 1.5 compared to vehicle controls (Fig. 7b and f). Furthermore, a significant treatment effect on wake bout duration (F_(1, 10)_ = 5.769, P=0.0372) was found showing that 2-AG treatment increased bout duration compared to vehicle controls in males (10.99 ± 0.7833 vs -31.47 ± 11.66) (Fig. 7c). Lastly, REMS was significantly decreased in 2-AG (-1.840 [-3.531, -0.1496] vs 1.640 [0.5208, 2.760]) treated male mice compared to vehicle controls over the 24-hour period (Treatment: F_(1, 10)_ = 12.56, P=0.0053; Time: F_(3.460, 34.60)_ = 3.616, P=0.0183). 2-AG treated male mice were slightly below baseline levels while vehicle controls were slightly above baseline levels. Post-hoc analysis following a significant interaction effect (F_(47, 470)_ = 2.033, P=0.0001) indicated that REMS was significantly depressed at ZTs 9.5 (-9.445 ± 2.03 vs 3.28 ± 1.42, P = 0.0292) and 12 (-9.83 ± 1.57 vs 4.30 ± 1.21, P=0.0021) from 2-AG treatment compared to vehicle (Fig. 7j). Lastly, analysis of REMS bout duration showed a significant main effect of treatment (F_(1, 10)_ = 10.55, P=0.0088) and time (F_(1, 10)_ = 1279, P<0.0001) in male mice revealing that 2-AG significantly decreased REMS bout duration compared to vehicle controls (-18.69 ± 16.73 vs -24.07 ± 12.22). Post-hoc testing following a significant interaction effect (F_(1, 10)_ = 31.02, P=0.0002) showed that the effect was restricted to the dark period (-1.96 ± 0.65 vs -11.85 ± 1.51, P<0.0001) (Fig. 7l).

**Fig. 7:**
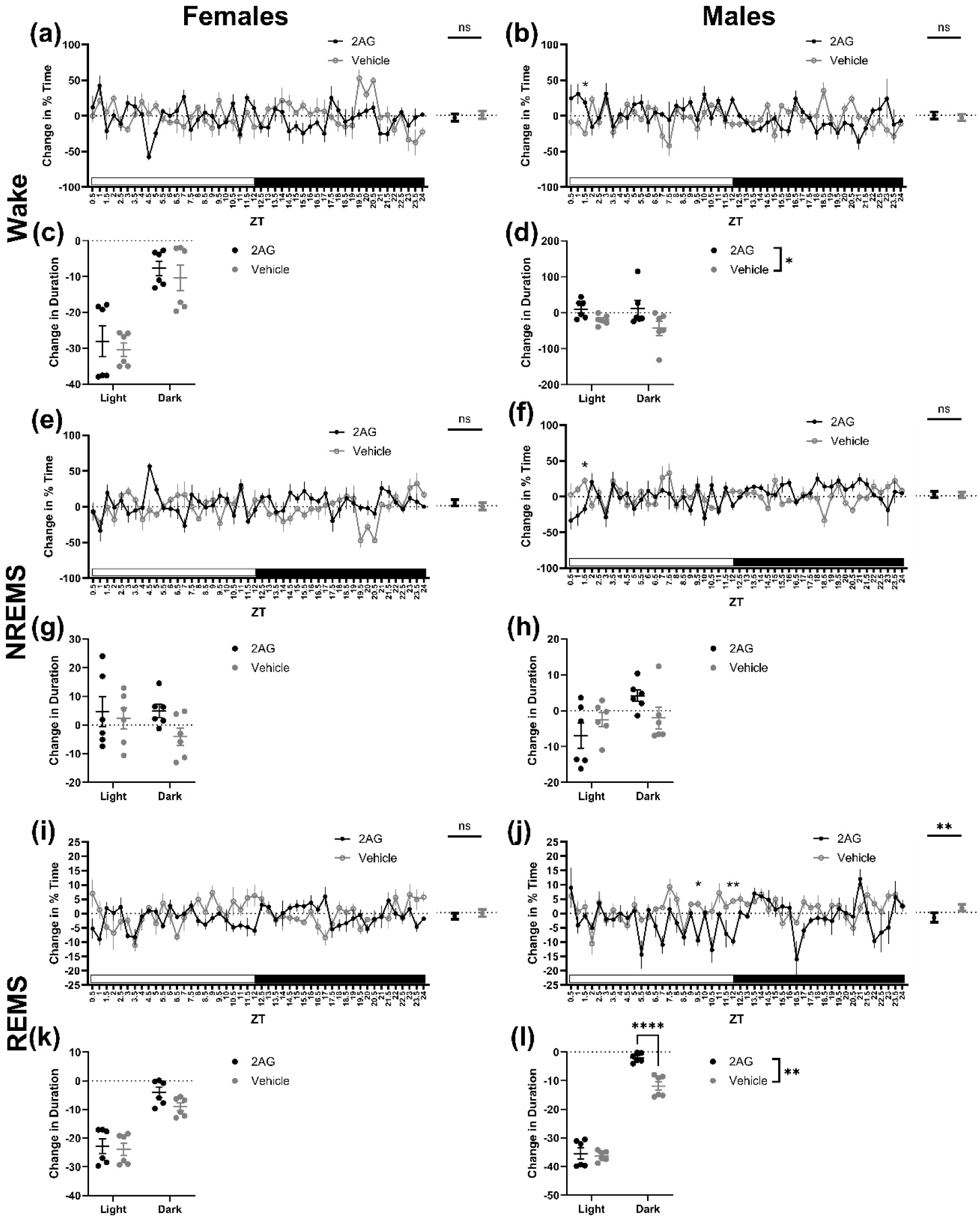
Sleep-wake changes in male and female mice during withdrawal day 3. (a) Half hourly (left) and total (right) change in percent time spent awake from baseline in females. (b) Half hourly (left) and total (right) change in percent time spent awake from baseline in males. (c) Change in wake bout duration from baseline in females during the light and dark period. (d) Change in wake bout duration from baseline in males during the light and dark period. (e) Half hourly (left) and total (right) change in percent time in NREMS from baseline in females. (f) Half hourly (left) and total (right) change in percent time in NREMS from baseline in males. (g) Change in NREMS bout duration from baseline in females during the light and dark period. (h) Change in NREMS bout duration from baseline in males during the light and dark period. (i) Half hourly (left) and total (right) change in percent time in NREMS from baseline in females. (j) Half hourly (left) and total (right) change in percent time in NREMS from baseline in males. (k) Change in REMS bout duration from baseline in females during the light and dark period. (l) Change in REMS bout duration from baseline in males during the light and dark period. Data presented as mean ± SEM for half hourly data and mean ± 95% CI for total data. ns = P > 0.05, * = P ≤ 0.05, ** = P ≤ 0.01, *** = P ≤ 0.001, **** = P ≤ 0.0001

## 4. Discussion

Sleep and circadian rhythm disruptions are common among people with OUD or opioid dependence. Fentanyl has become the most commonly misused synthetic opioid. The exact nature of sleep disruption through a cycle of fentanyl addiction remains unknown. Importantly, while effective opioid substitution medications exist (e.g., methadone), there are currently limited non-opioid based drugs that can be used to mitigate withdrawal symptoms including sleep disruption. We characterized diurnal activity and sleep-wake changes from fentanyl injection administration. During withdrawal from fentanyl, we examined whether the prominent endocannabinoid 2-AG could improve sleep during fentanyl-induced withdrawal in male and female mice.

We show that chronic fentanyl treatment decreased wake while increasing NREMS and REMS. While no detailed sleep-wake data for chronic fentanyl use in humans exists, chronic treatment of other opioids has been indicated to increase light NREMS while depressing deep NREMS (Wang and Teichtahl, 2007). Given that light NREMS makes up the majority of total NREMS time (Carskadon and Dement, 2011) in humans, our data are likely in agreement with the human data. The piezo sensor leveraged in this study, however, cannot separate the stages of NREMS and further research will be necessary to confirm this effect. Moreover, we show that REMS increased following injections compared to baseline with days 1 through 3 being the most robust. This effect diminished over time, yet no statistical difference was detected in REMS between day 1 and day 7 in either sex. Most studies in humans indicate that various opioids lead to reduced REM;, however, the sample sizes in these studies are exceedingly low (most under 10 individuals), only a few include females, and most have no controls (Wang and Teichtahl, 2007). The increase in REMS seen here may be fentanyl or mouse specific, but also highlights that the effects of opioids on REMS remain uncertain.

Our previous work showed that NREMS progressively decreased during withdrawal (and wake in turn increased) compared to baseline levels (Gamble et al., 2022). Here we showed that during spontaneous withdrawal mice treated with DMSO, drug vehicle, displayed increased or unchanged NREMS (and decreased wakefulness) compared to baseline. DMSO has been shown to increase light NREMS starting at concentrations of 15% in male rats (Cavas et al., 2005). While we dissolved 2-AG in 10% DMSO deliberately to avoid this effect, the minimum sleep potentiating dose in mice is unknown for either sex. Furthermore, DMSO has shown to have numerous physiological effects including when administered with morphine such as altered antinociception (Fossum et al., 2008) and potentially increased sleep (Caselli et al., 2009) which might similarly alter the effects of opioid-withdrawal.

Overall, 2-AG treatment produced the opposite effect of vehicle DMSO treatment in withdrawal days 1 through 3. While 2-AG often restored sleep-wake levels to baseline, given the sleep promoting effects of DMSO it is conceivable that 2-AG treatment may make withdrawal worse causing more wakefulness and less NREMS. Indeed, while 2-AG increased arousal and wake bout duration on withdrawal day 1 in both sexes and withdrawal day 2 in females only, this effect was mostly confined to the dark (active) period. This suggests a diurnal effect of 2-AG treatment and a potential value in addressing sleepiness. Interestingly, by withdrawal day 3, 2-AG treatment caused a notable decrease in REMS compared to baseline (and controls) in males only. REMS bout duration in males was also preserved at baseline levels was also preserved at baseline levels in dark, but not light period. The literature on the effects of opioids on REMS during withdrawal is sparse but suggest increased REMS during withdrawal from heroin or methadone in male individuals (Wang and Teichtahl, 2007). Previous work from our lab using electroencephalography (EEG) in male mice detected no changes in REMS on the final day of chronic fentanyl injections at the same dose. Work in rats has shown that dronabinol decreases REMS in a CB receptor independent manner (Calik and Carley, 2017). Interestingly, Calik and Carley show that this effect is not DMSO-dependent, but that DMSO does in part enhance the effect of dronabinol on sleep (Calik and Carley, 2019). With respect to wake and NREMS levels during withdrawal day 3, only wake bouts were slightly elevated to baseline in 2-AG compared to DMSO in males while all other measures were unchanged.

There are several limitations of the present work. First, no non-drug injection control was included so it is not possible to tease out what differences in sleep-wake during administration and withdrawal are only due to fentanyl and 2-AG, respectively. Second, piezo sleep recording relies on breathing to determine REMS classification which is altered by fentanyl (Hill et al., 2020). This effect has been noted to last ∼3 hours when morphine is administered (Kamei et al., 2011) and may partially explain the increased REMS after the first injection in days 1 through 3. This effect presumably decreases as tolerance develops. To confirm these findings, a detailed follow-up study using EEG is warranted.

While unclear that 2-AG can ameliorate NREMS disruption due to opioid withdrawal, it may be able to combat sleepiness induced by sleep disruption by facilitating wakefulness and stabilize REMS. Still, replication and further investigation as to the exact mechanism and is needed in order to establish any therapeutic potential of targeting the eCB system to treat opioid withdrawal.

## Funding

Work was supported by National Heart, Lung, and Blood Institute (NHLBI) R01HL150432 (RWL) funded by the National Institutes of Health (NIH) Helping End Addiction Long-term (HEAL) Initiative.

## Author Contributions

**R.W.L., M.C.G., and S.M.:** Conceptualization, **M.C.G. and S.M.:** Data curation, **R.W.L**., **M.C.G., and S.M.:** Formal analysis, **R.W.L:** Funding acquisition, **M.C.G. and S.M.:** Investigation, **R.W.L., M.C.G., and S.M.:** Methodology, **R.W.L.** and **M.C.G.:** Project administration, **R.W.L:** Supervision, **M.C.G., and S.M.:** Visualization, **R.W.L., M.C.G., S.M., and B.R.W:** Roles/Writing - original draft, **R.W.L., M.C.G., S.M., and B.R.W:** Writing - review & editing.

## Notes

### Competing Interest Statement

The authors have declared no competing interest.

## References

Ahmad FB, Cisewski JA, Rossen LM, Sutton P, 2023. Provisional drug overdose death counts [WWW Document]. URL https://www.cdc.gov/nchs/nvss/vsrr/drug-overdose-data.htm (accessed 5.20.23).

Bird, H.E., Huhn, A.S., Dunn, K.E., 2023. Fentanyl Absorption, Distribution, Metabolism, and Excretion: Narrative Review and Clinical Significance Related to Illicitly Manufactured Fentanyl. J Addict Med Publish Ahead of Print. 10.1097/ADM.0000000000001185

Caillé, S., Alvarez-Jaimes, L., Polis, I., Stouffer, D.G., Parsons, L.H., 2007. Specific Alterations of Extracellular Endocannabinoid Levels in the Nucleus Accumbens by Ethanol, Heroin, and Cocaine Self-Administration. J. Neurosci. 27, 3695–3702. 10.1523/JNEUROSCI.4403-06.2007

Calik, M.W., Carley, D.W., 2019. DMSO potentiates the suppressive effect of dronabinol, a cannabinoid, on sleep apnea and REM sleep (preprint). Neuroscience. 10.1101/769463

Calik, M.W., Carley, D.W., 2017. Effects of Cannabinoid Agonists and Antagonists on Sleep and Breathing in Sprague-Dawley Rats. Sleep 40, zsx112. 10.1093/sleep/zsx112

Carskadon, M.A., Dement, W.C., 2011. Chapter 2 – Normal Human Sleepc: An Overview. Principles and practice of sleep medicine.

Caselli, D., Tintori, V., Messeri, A., Frenos, S., Bambi, F., Aricò, M., 2009. Respiratory depression and somnolence in children receiving dimethylsulfoxide and morphine during hematopoietic stem cells transplantation. Haematologica 94, 152–153. 10.3324/haematol.13828

Cavas, M., Beltrán, D., Navarro, J.F., 2005. Behavioural effects of dimethyl sulfoxide (DMSO): Changes in sleep architecture in rats. Toxicology Letters 157, 221–232. 10.1016/j.toxlet.2005.02.003

Chieng, B., Williams, J.T., 1998. Increased Opioid Inhibition of GABA Release in Nucleus Accumbens during Morphine Withdrawal. J. Neurosci. 18, 7033–7039. 10.1523/JNEUROSCI.18-17-07033.1998

Fossum, E.N., Lisowski, M.J., Macey, T.A., Ingram, S.L., Morgan, M.M., 2008. Microinjection of the vehicle dimethyl sulfoxide (DMSO) into the periaqueductal gray modulates morphine antinociception. Brain Research 1204, 53–58. 10.1016/j.brainres.2008.02.022

Galaj, E., Xi, Z.-X., 2019. Potential of Cannabinoid Receptor Ligands as Treatment for Substance Use Disorders. CNS Drugs 33, 1001–1030. 10.1007/s40263-019-00664-w

Gamble, M.C., Chuan, B., Gallego-Martin, T., Shelton, M.A., Puig, S., O’Donnell, C.P., Logan, R.W., 2022. A role for the circadian transcription factor NPAS2 in the progressive loss of non-rapid eye movement sleep and increased arousal during fentanyl withdrawal in male mice. Psychopharmacology. 10.1007/s00213-022-06200-x

Hanlon, E.C., Tasali, E., Leproult, R., Stuhr, K.L., Doncheck, E., de Wit, H., Hillard, C.J., Van Cauter, E., 2016. Sleep Restriction Enhances the Daily Rhythm of Circulating Levels of Endocannabinoid 2-Arachidonoylglycerol. Sleep 39, 653–664. 10.5665/sleep.5546

Hill, R., Santhakumar, R., Dewey, W., Kelly, E., Henderson, G., 2020. Fentanyl depression of respiration: Comparison with heroin and morphine. British Journal of Pharmacology 177, 254–265. 10.1111/bph.14860

Kamei, J., Ohsawa, M., Hayashi, S.-S., Nakanishi, Y., 2011. Effect of chronic pain on morphine-induced respiratory depression in mice. Neuroscience 174, 224–233. 10.1016/j.neuroscience.2010.11.012

Kay, D.C., 1975. Human sleep during chronic morphine intoxication. Psychopharmacologia 44, 117–124. 10.1007/BF00420997

Kendall, D.A., Yudowski, G.A., 2017. Cannabinoid Receptors in the Central Nervous System: Their Signaling and Roles in Disease. Frontiers in Cellular Neuroscience 10.

Kesner, A.J., Lovinger, D.M., 2020. Cannabinoids, Endocannabinoids and Sleep. Front Mol Neurosci 13, 125. 10.3389/fnmol.2020.00125

Lewis, S.A., Oswald, I., Evans, J.I., Akindele, M.O., Tompsett, S.L., 1970. Heroin and human sleep. Electroencephalography and Clinical Neurophysiology 28, 374–381. 10.1016/0013-4694(70)90230-0

Martin, M., Ledent, C., Parmentier, M., Maldonado, R., Valverde, O., 2000. Cocaine, but not morphine, induces conditioned place preference and sensitization to locomotor responses in CB1 knockout mice. European Journal of Neuroscience 12, 4038– 4046. 10.1046/j.1460-9568.2000.00287.x

Nathan, P.J., O’Neill, B.V., Napolitano, A., Bullmore, E.T., 2011. Neuropsychiatric Adverse Effects of Centrally Acting Antiobesity Drugs. CNS Neuroscience & Therapeutics 17, 490–505. 10.1111/j.1755-5949.2010.00172.x

Navarro, M., Carrera, M.R.A., del Arco, I., Trigo, J.M., Koob, G.F., Rodríguez de Fonseca, F., 2004. Cannabinoid receptor antagonist reduces heroin self-administration only in dependent rats. European Journal of Pharmacology 501, 235–237. 10.1016/j.ejphar.2004.08.022

Neumueller, S.E., Buiter, N., Hilbert, G., Grams, K., Taylor, R., Desalvo, J., Hodges, G.L., Hodges, M.M., Pan, L.G., Lewis, S.J., Forster, H.V., Hodges, M.R., 2023. Effects of sub-lethal doses of fentanyl on vital physiologic functions and withdrawal-like behaviors in adult goats. Frontiers in Physiology 14.

Oswald, I., 1969. Sleep, dreaming and drugs. Proc R Soc Med 62, 151–153.

Pava, M.J., Hartog, C.R. den, Blanco-Centurion, C., Shiromani, P.J., Woodward, J.J., 2014. Endocannabinoid Modulation of Cortical Up-States and NREM Sleep. PLOS ONE 9, e88672. 10.1371/journal.pone.0088672

Pava, M.J., Makriyannis, A., Lovinger, D.M., 2016. Endocannabinoid Signaling Regulates Sleep Stability. PLoS One 11, e0152473. 10.1371/journal.pone.0152473

Pérez-Morales, M., De La Herrán-Arita, A.K., Méndez-Díaz, M., Ruiz-Contreras, A.E., Drucker-Colín, R., Prospéro-García, O., 2013. 2-AG into the lateral hypothalamus increases REM sleep and cFos expression in melanin concentrating hormone neurons in rats. Pharmacology Biochemistry and Behavior 108, 1–7. 10.1016/j.pbb.2013.04.006

Pickel, V.M., Chan, J., Kash, T.L., Rodríguez, J.J., MacKie, K., 2004. Compartment-specific localization of cannabinoid 1 (CB1) and μ-opioid receptors in rat nucleus accumbens. Neuroscience 127, 101–112. 10.1016/j.neuroscience.2004.05.015

Puig, S., Shelton, M.A., Barko, K., Seney, M.L., Logan, R.W., 2022. Sexcspecific role of the circadian transcription factor NPAS2 in opioid tolerance, withdrawal and analgesia. Genes Brain Behav 21, e12829. 10.1111/gbb.12829

Ramesh, D., Gamage, T.F., Vanuytsel, T., Owens, R.A., Abdullah, R.A., Niphakis, M.J., Shea-Donohue, T., Cravatt, B.F., Lichtman, A.H., 2013. Dual Inhibition of Endocannabinoid Catabolic Enzymes Produces Enhanced Antiwithdrawal Effects in Morphine-Dependent Mice. Neuropsychopharmacol 38, 1039–1049. 10.1038/npp.2012.269

Ramesh, D., Ross, G.R., Schlosburg, J.E., Owens, R.A., Abdullah, R.A., Kinsey, S.G., Long, J.Z., Nomura, D.K., Sim-Selley, L.J., Cravatt, B.F., Akbarali, H.I., Lichtman, A.H., 2011. Blockade of Endocannabinoid Hydrolytic Enzymes Attenuates Precipitated Opioid Withdrawal Symptoms in Mice. J Pharmacol Exp Ther 339, 173–185. 10.1124/jpet.111.181370

Saigusa, T., Aono, Y., Waddington, J.L., 2021. Integrative opioid-GABAergic neuronal mechanisms regulating dopamine efflux in the nucleus accumbens of freely moving animals. Pharmacol. Rep 73, 971–983. 10.1007/s43440-021-00249-9

Shah, M., Huecker, M.R., 2023. Opioid Withdrawal, in: StatPearls [Internet]. StatPearls Publishing.

Sloan, M.E., Gowin, J.L., Ramchandani, V.A., Hurd, Y.L., Le Foll, B., 2017. The endocannabinoid system as a target for addiction treatment: Trials and tribulations. Neuropharmacology, A New Dawn in Cannabinoid Neurobiology 124, 73–83. 10.1016/j.neuropharm.2017.05.031

Sugiura, T., Kishimoto, S., Oka, S., Gokoh, M., 2006. Biochemistry, pharmacology and physiology of 2-arachidonoylglycerol, an endogenous cannabinoid receptor ligand. Progress in Lipid Research 45, 405–446. 10.1016/j.plipres.2006.03.003

Sustkova-Fiserova, M., Charalambous, C., Havlickova, T., Lapka, M., Jerabek, P., Puskina, N., Syslova, K., 2017. Alterations in Rat Accumbens Endocannabinoid and GABA Content during Fentanyl Treatment: The Role of Ghrelin. Int J Mol Sci 18, 2486. 10.3390/ijms18112486

Valverde, O., Noble, F., Beslot, F., Daugé, V., Fournié-Zaluski, M.-C., Roques, B.P., 2001. Δ9-tetrahydrocannabinol releases and facilitates the effects of endogenous enkephalins: reduction in morphine withdrawal syndrome without change in rewarding effect. European Journal of Neuroscience 13, 1816–1824. 10.1046/j.0953-816x.2001.01558.x

Viganò, D., Grazia Cascio, M., Rubino, T., Fezza, F., Vaccani, A., Di Marzo, V., Parolaro, D., 2003. Chronic Morphine Modulates the Contents of the Endocannabinoid, 2-Arachidonoyl Glycerol, in Rat Brain. Neuropsychopharmacol 28, 1160–1167. 10.1038/sj.npp.1300117

Wang, D., Teichtahl, H., 2007. Opioids, sleep architecture and sleep-disordered breathing. Sleep Medicine Reviews 11, 35–46. 10.1016/j.smrv.2006.03.006

Wenzel, J.M., Cheer, J.F., 2018. Endocannabinoid Regulation of Reward and Reinforcement through Interaction with Dopamine and Endogenous Opioid Signaling. Neuropsychopharmacol. 43, 103–115. 10.1038/npp.2017.126

Xi, Z.-X., Stein, E.A., 2002. GABAergic MECHANISMS OF OPIATE REINFORCEMENT. Alcohol and Alcoholism 37, 485–494. 10.1093/alcalc/37.5.485

Yamaguchi, T., Hagiwara, Y., Tanaka, H., Sugiura, T., Waku, K., Shoyama, Y., Watanabe, S., Yamamoto, T., 2001. Endogenous cannabinoid, 2-arachidonoylglycerol, attenuates naloxone-precipitated withdrawal signs in morphine-dependent mice. Brain Research 909, 121–126. 10.1016/S0006-8993(01)02655-5

